# Elasticity of dense actin networks produces nanonewton protrusive forces

**DOI:** 10.1101/2021.04.13.439622

**Authors:** Marion Jasnin, Jordan Hervy, Stéphanie Balor, Anais Bouissou, Amsha Proag, Raphaël Voituriez, Isabelle Maridonneau-Parini, Wolfgang Baumeister, Serge Dmitrieff, Renaud Poincloux

## Abstract

Actin filaments assemble into force-generating systems involved in diverse cellular functions, including cell motility, adhesion, contractility and division. It remains unclear how networks of actin filaments, which individually generate piconewton forces, can produce forces reaching tens of nanonewtons. Here we use *in situ* cryo-electron tomography to unveil how the nanoscale architecture of macrophage podosomes enables basal membrane protrusion. We show that the sum of the actin polymerization forces at the membrane is not sufficient to explain podosome protrusive forces. Quantitative analysis of podosome organization demonstrates that the core is composed of a dense network of bent actin filaments storing elastic energy. Theoretical modelling of the network as a spring-loaded elastic material reveals that it exerts forces of up to tens of nanonewtons, similar to those evaluated experimentally. Thus, taking into account not only the interface with the membrane but also the bulk of the network, is crucial to understand force generation by actin machineries. Our integrative approach sheds light on the elastic behavior of dense actin networks and opens new avenues to understand force production inside cells.

## Introduction

Actin, one of the most abundant proteins in eukaryotic cells, organize into force-generating filamentous networks, which play pivotal roles in cell motility, adhesion, endocytosis and vesicular traffic ^1,2^. Thermodynamics showed that actin polymerization generates mechanical forces by the addition of new monomers at the end of a fluctuating filament ^3,4^ Typical stall forces of a polymerizing actin filament were estimated in the 1-10 pN range ^3,5,6^, in agreement with experimental values of 1.5 pN found by optical trap measurements ^7^. Thus, polymerization of actin filaments against a membrane is capable of extruding thin membrane tubes or forming plasma membrane invaginations in mammalian cells in a force range of a few tens of pN ^8–10^.

The forces exerted by actin filaments can also reach a much higher regime, in the nanonewton (nN) range ^11,12^. At the leading edge of motile cells, branched actin networks in lamellipodial protrusions produce local forces of ~1 nN ^13,14^. During yeast endocytosis, the actin machinery generates similar forces to overcome the turgor pressure pushing the invagination outwards ^15^. At the basal membrane of myeloid cells, podosomes probe the stiffness of the extracellular environment through the generation of forces reaching tens of nN ^16^. Unlike the lower force regime, the mechanisms by which meshworks of actin filaments produce nN forces remain unknown.

If the force generated by an actin network corresponds to the sum of the polymerization forces generated by single filaments pushing against a load, then hundreds to thousands of filaments would need to continuously grow against the surface to reach the nN range. Alternatively, filaments could push on the membrane with shallow angles ^17,18^. It was also proposed that the actin network could store elastic energy, and thus exert a restoring force. In the low force regime, bending of endocytic filaments observed in animal cells has been proposed to store elastic energy for pit internalization ^19^. In the large force regime, a theoretical model with non-deformable filaments showed that a large force would cause elastic energy to be stored in cross-linkers deformation ^20^. However, none of these hypotheses have been explored in native actin machineries generating forces of several nN, due to the difficulty to combine direct observation of the actin network architecture and the knowledge of the exerted force.

To date, cryo-electron tomography (cryo-ET) is the only technique that resolves single actin filaments inside unperturbed cellular environments ^21–23^. Here we use cryo-ET to unveil the three-dimensional (3D) architecture of human macrophage podosomes and elucidate their force generation mechanism. These submicrometric structures are composed of a protrusive core of actin filaments surrounded by an adhesion ring. The balance of forces requires the protrusion force applied by the core on the substrate to be counteracted by a force of equal magnitude. Protrusion force microscopy (PFM) revealed that this balance of forces takes place locally through traction at the adhesion ring ^16,24,25^, which has been proposed to be transmitted by radial actin cables connecting the core to the ring ^25^. Here we visualize these radial filaments and quantitatively analyze the 3D organization of the core and ring networks. We show that the protrusive forces generated by podosomes cannot be explained by the sum of the actin polymerization forces at the core membrane but by the storage of elastic energy into the dense network of bent actin filaments. These results explain how cellular actin networks can act as a spring-loaded elastic material to exert forces of up to tens of nanonewtons.

## Results

### Cryo-ET allows quantitative analysis of filament organization in podosomes

Owing to their size and location at the basal cell membrane, native podosomes are amenable to cryo-ET exploration using cryo-focused ion beam (cryo-FIB) milling sample preparation. We prepared thin vitrified sections (so-called wedges) containing podosomes using shallow incidence angles of the ion beam and subjected them to cryo-ET (Figure 1A and Movie S1). Segmentation of the tomograms revealed that podosomes are made of a core of oblique filaments surrounded by radial filaments (Figure 1B-C and Movie S1), as predicted previously ^25^. Other cytoskeletal elements, cellular organelles, ribosomes and glycogen granules are excluded from the core and radial actin networks, gathering either at the periphery or on top of podosomes (Figure 1A-B and Movie S1).

**Fig. 1:**
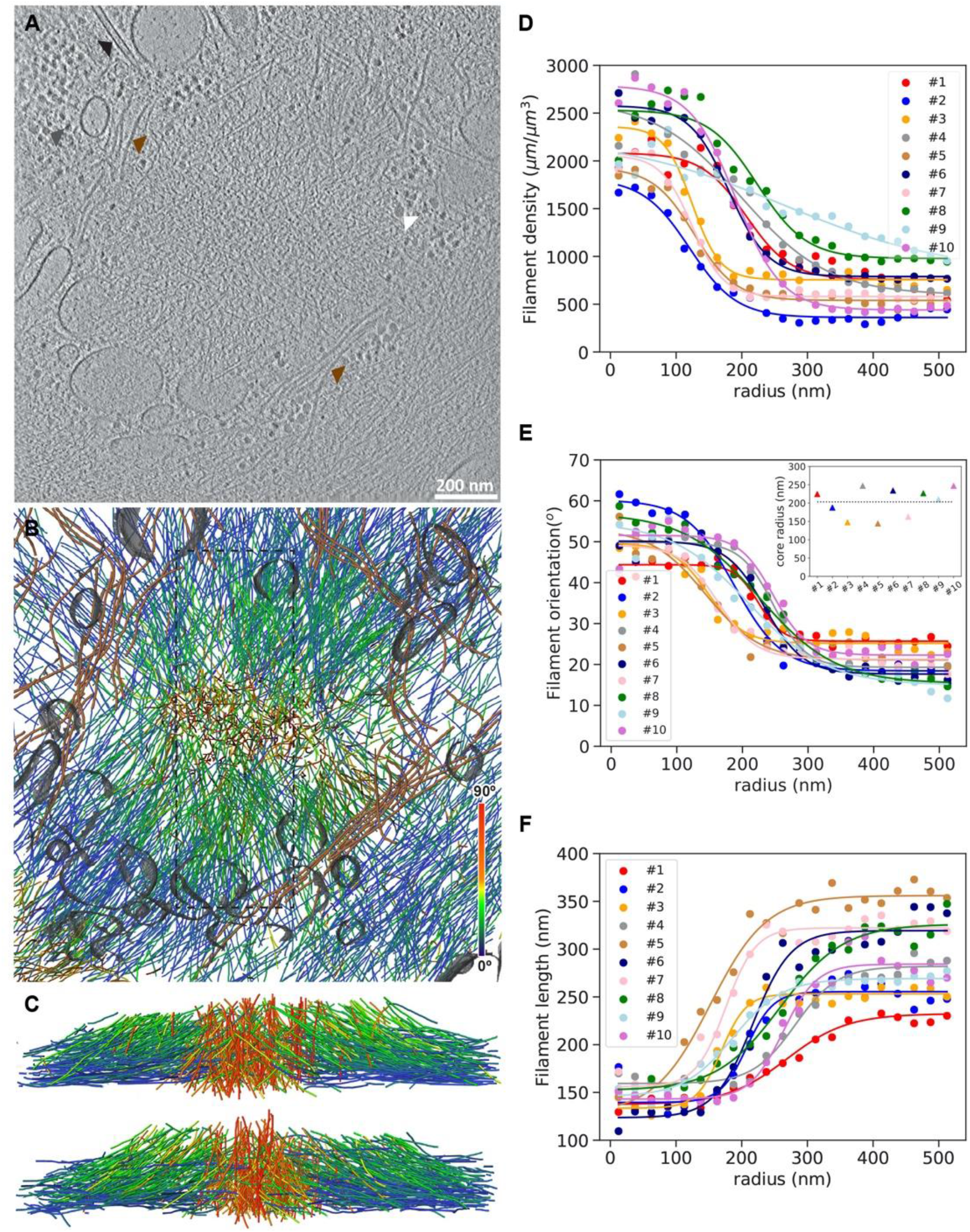
Cryo-electron tomography allows quantitative analysis of actin filament organization in native podosomes. **A:** Slice from a tomographic volume acquired in a frozen-hydrated human macrophage revealing the podosome environment. Colored arrows point to ribosomes (grey), glycogen granules (white), a microtubule (black), and intermediate filaments (brown). See also Movie S1. **B:** Orthographic view of the corresponding 3D segmentation of the actin filaments showing their relative orientation with respect to the basal membrane, intermediate filaments (brown), and organelle membranes (grey). **C:** Perspective views of the actin filaments in the volume indicated by a dotted rectangle in (B) shown from the left (top) and right (bottom) sides. **D-F:** Average filament density (**D**), orientation (**E**) and length (**F**) as a function of the radial distance from the core center for ten tomograms. Inset in (**E**): Corresponding values estimated for the core radius (Methods). The podosome shown in (**A-C**) corresponds to #9.

Quantitative analysis of filament organization highlighted the specificity of the actin core in terms of filament length, density and orientation with respect to the basal membrane (Methods and Figures S1–S3). All of these parameters exhibit a sharp transition as a function of the radial distance from the core center (Figure 1D-F). Density is higher inside the core by a factor of 2 to 3 (Figure 1D) and filaments display a mean orientation of 5O±21 °relative to the plasma membrane, as compared to the flatter 23±21°outside the core (Figure 1E and Figure S4A-B). Core filaments are shorter than the surrounding radial filaments, with mean lengths of 119±52 nm and 181±134 nm, respectively (Figure 1F and Figure S4C-D). Further analysis of the transition curves for the filament orientation provided a mean core radius of 203±38 nm for a total of ten podosomes (Figure 1E, inset). This agrees with the values obtained from the fits of the other parameters (Figure S5).

Since the milling procedure removed the top of the podosomes, we also imaged podosomes exposed by cell unroofing prior to cryo-fixation to get a complete picture of podosome organization (Figure S6 and Movie S2). In addition to the core and radial filaments observed previously, we detected horizontal filaments on top of the core as well as in between neighboring cores. These filaments, which may have been lifted up from the plasma membrane during podosome growth, could also participate in the generation of the traction forces that counterbalance the protrusion forces generated by podosomes ^25^.

### The sum of the actin polymerization forces at the core membrane is below 1 nN

To evaluate the forces generated by actin polymerization on the plasma membrane beneath the podosome core, where protrusion occurs ^16,24,25^, we identified filament segments in the close vicinity of and protruding against the plasma membrane (Figure 2A and Movie S3). We found an average of 45±29 filaments per podosome, with a mean orientation of 61±6°relative to the plasma membrane (Figure 2B). This rules out the first two hypotheses, namely that 1) hundreds to thousands of filaments are growing at the same time onto the plasma membrane, or that 2) filaments are pushing with shallow angles. In addition, using the upper limit of 10 pN for the stall force of a single filament growing perpendicularly to the membrane ^3,5,6,26^, taking into account filament orientation ^17^, and considering that all these filaments are polymerizing concomitantly, we evaluated a maximal polymerization force of 615±396 pN per podosome (ten podosomes evaluated; Methods and Figure 2C-D). This is one order of magnitude lower than the experimental values ^16^. We therefore concluded that the total force produced by polymerization of actin filaments against the plasma membrane is not sufficient to generate the protrusive forces exerted by podosomes.

**Fig. 2.**
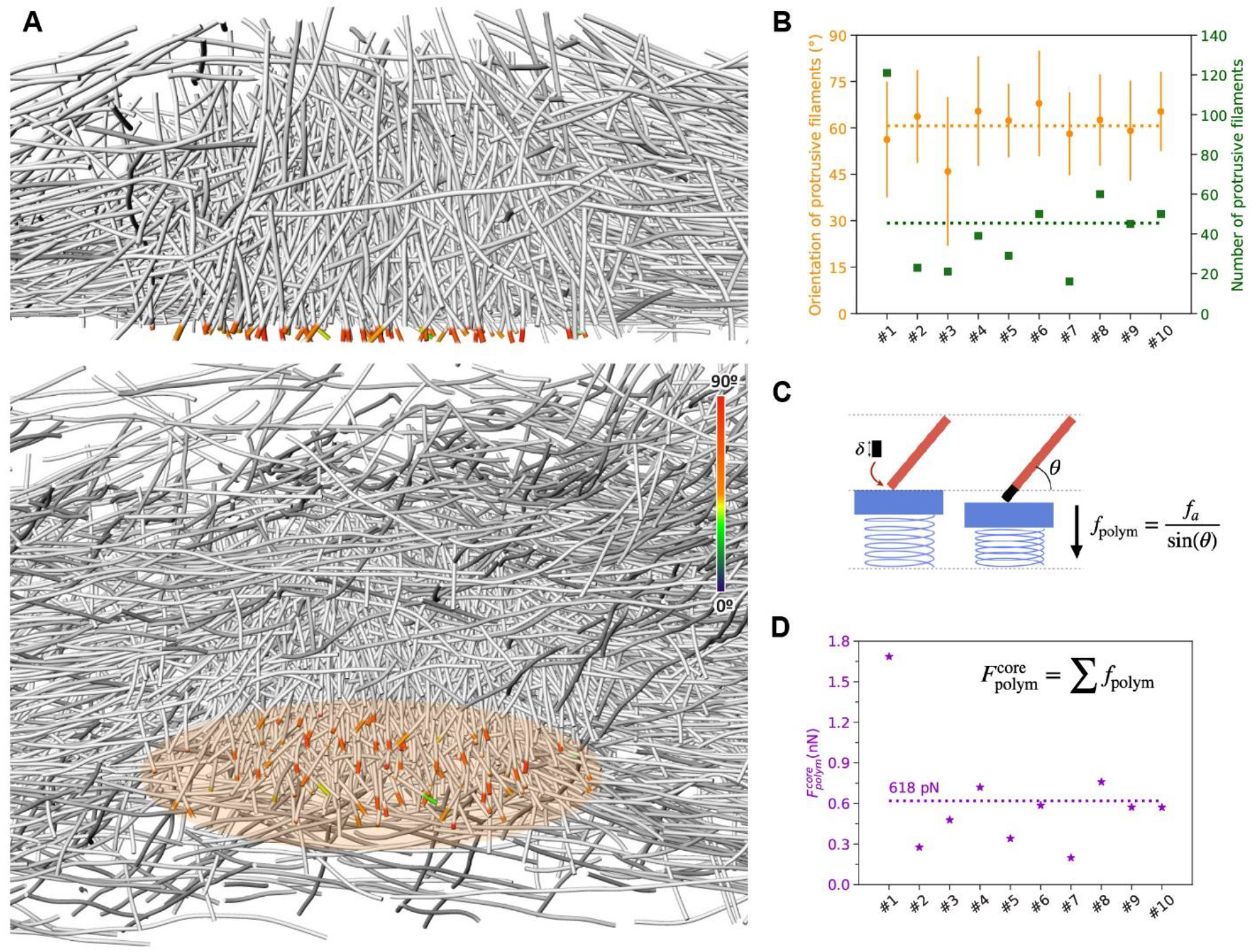
The sum of the actin polymerization forces at the core membrane is below 1 nN. **A:** (Top) Perspective view of a cross-section through podosome #8 displaying in color the part of the core filaments pushing against the plasma membrane. (Bottom) Orthographic view of the same podosome from a different angle. The color map corresponds to the filament orientation with respect to the basal membrane. See also Movie S3. **B:** Mean orientation (orange) and number (green) of polymerizing filaments at the core membrane. **C:** Scheme representing the polymerization force generated by the addition of a new monomer at the growing end of a filament with an inclination *θ* relative to the membrane. **D:** Estimated polymerization force generated at the core membrane.

### The actin core stores elastic energy

Next, we tested the third hypothesis, that is, the storage of elastic energy in the network. Visual inspection of the networks indicated that podosome filaments are bent (Figure 3A and Movie S4). The projection of each filament on the vertical plane passing through its two ends highlighted the variation of their local curvature along the filament length (Methods and Figure 3B-C). Quantitative analysis of filament curvature revealed an average compressive strain of 4.2 ±0.4% for all filaments, independently of their localization within the podosome (Figure 3D and Figure S7). We therefore evaluated the elastic energy using the theory of linear elasticity: actin filaments can be modeled as semi-flexible polymers with the following elastic energy at the single filament level ^27^:

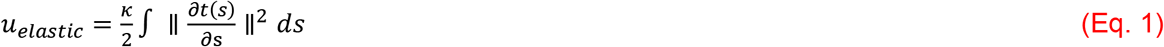

where *k* ~ 4*10^-26^ N.m^2^ is the bending modulus ^28^, *t*(*s*) is the tangent vector as a function of the arc-length coordinate *s*, and the integrand represents the square of the local curvature along the filament (Methods and Figure S8A). We evaluated the elastic energy per unit volume as the sum of the energies over all filaments in a given interval at a radial distance *r* from the core (Figure S8B). The density of elastic energy stored inside the core is much larger than that outside the core, by factors ranging from 3 to 10 (Figure 3E). This is consistent with the larger actin density by factors of 2 to 6 measured inside the core (Figure 1D). Therefore, this specific architecture allows the system to store elastic energy inside the podosome core, ranging from ~10^4^*k_B_T* to ~5.10^4^*k_B_T*. This corresponds to 40*k_B_T* per filament on average, which is much larger than the scale of thermal fluctuations.

**Fig. 3.**
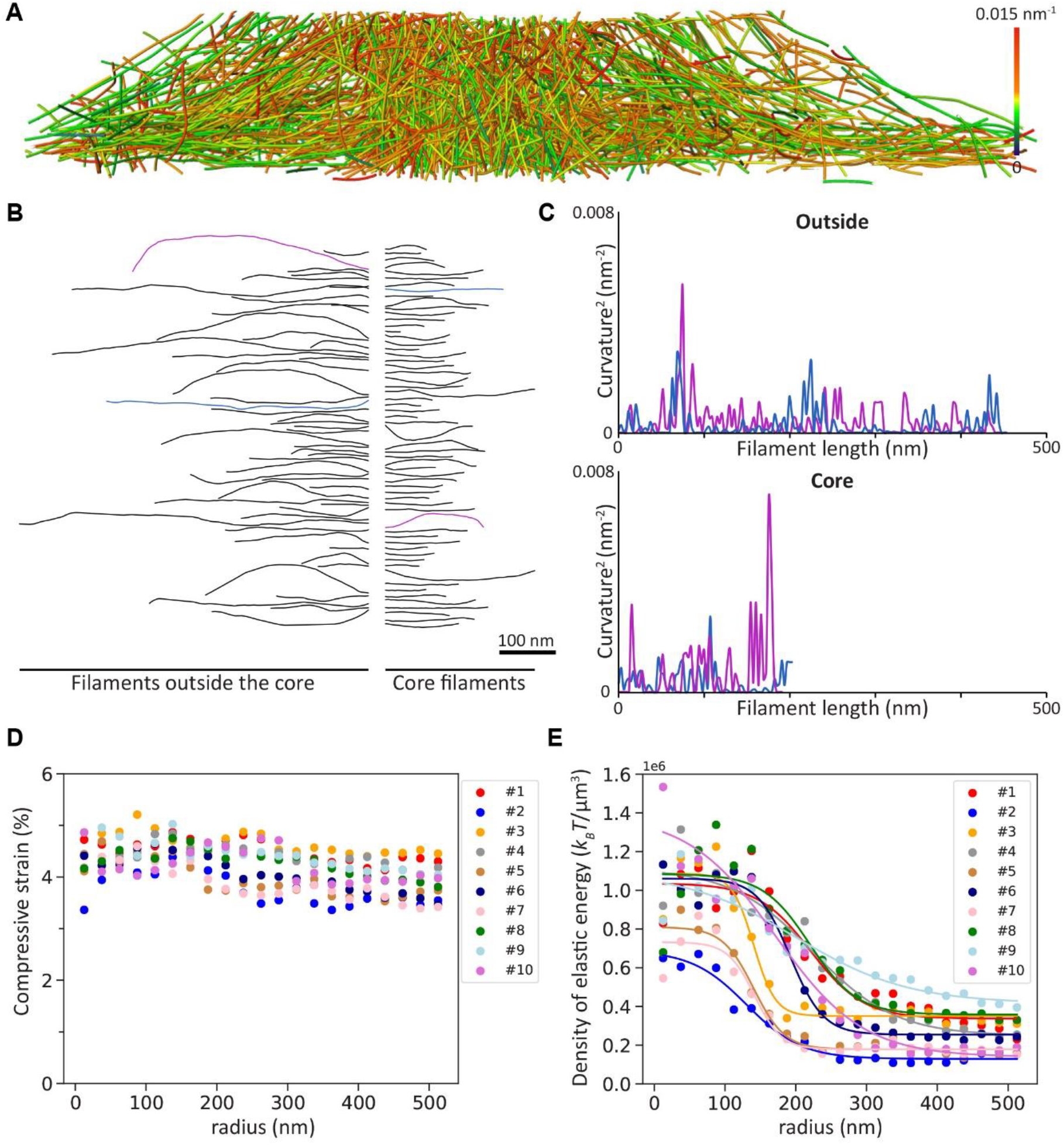
Podosome filaments are compressed and store high elastic energy inside the core. **A:** Perspective view of a cross-section through podosome #1 displaying the mean curvature of the filaments. See also Movie S4. **B:** Gallery of a random selection of actin filaments from podosome #1. Filaments from the core (right) and outside the core (left) are shown. **C:** Square of the local curvature along the filament length for the blue and purple filaments from the galleries of filaments outside (top) and inside (bottom) the core shown in (**B**). **D:** Average compressive strain as a function of the radial distance from the core center for ten tomograms. **E:** Density of elastic energy as a function of the radial distance from the core center for ten tomograms.

### The actin core generates an elastic force in the nN range

To test whether the stored elastic energy can account for podosome protrusion forces, we next evaluated the elastic force generated through the compression of the actin network. Assuming that the podosome core behaves as a homogeneous elastic material, the force exerted by the core is such that its work for a small deformation of amplitude *δh_core_* is equal to 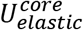 (the total elastic energy stored in the podosome core), with *δh_core_ = ϵ_core_ × h_core_* (*ε_core_* is the average filament compressive strain in the core and *h_core_* is the core height; Methods). We thus find:

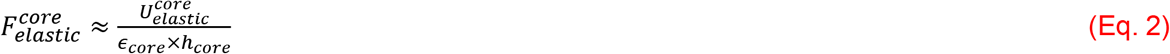

Using Eq. 2, we found an average elastic force of 9.7±4.2 nN, close to the mean value of 1O.4±3.8 nN reported from PFM measurements ^16^. The elastic force varies significantly between podosomes, with values ranging from 4.1 to 14.4 nN (Figure 4A). This can be explained by the disparity in the core size in our tomograms. Consistently with this, the force per unit area of the core exhibits less variation with an average value of *P* = 71.1 ± 12.9 kPa (Methods). This is in agreement, within the experimental margin of error, with the pressure estimated by PFM ^16^(Methods and Figure 4B) and in the same order of magnitude as that produced during endocytosis in yeast ^18^. Note that the elastic energy stored by the network, and thus the elastic force, are underestimated here since we do not take into account the elastic energy stored by crosslinkers^20^

**Fig. 4.**
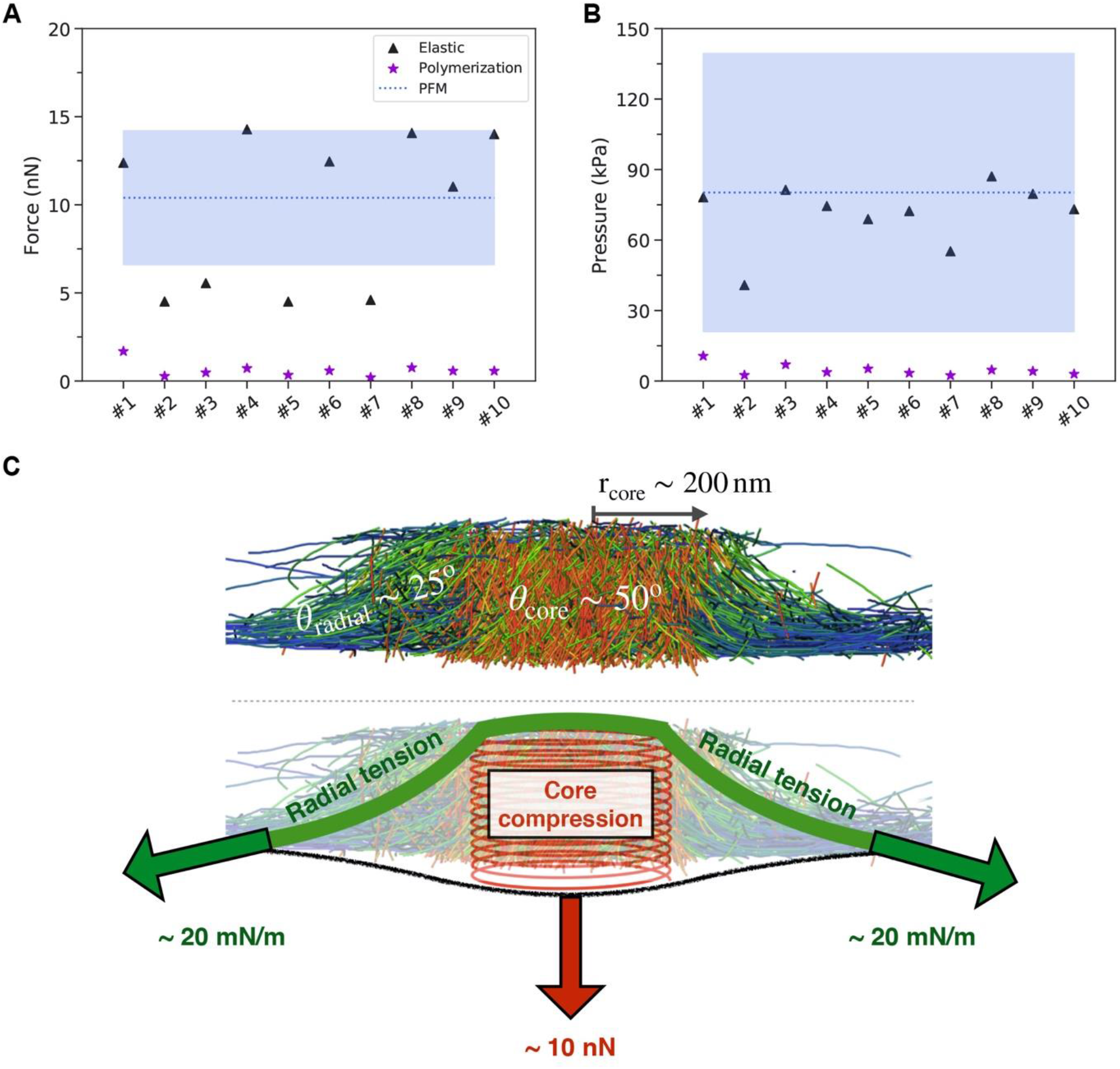
The actin core generates elastic forces in the nanonewton range. **A:** Comparison of the elastic force (“Elastic”) generated by the core through the compression of the actin network (black triangle) with the estimated polymerization force (“Polymerization”) generated at the core membrane (purple stars). The blue dashed line corresponds to the mean force derived from PFM measurements (“PFM”) ^16^ and the rectangular area filled in blue represents its standard deviation. **B:** Same comparison plot as in (**A**) for the estimated pressure assuming a perfect circular shape for the podosome core (Methods). **C:** Summary scheme showing the podosome organization revealed by cryo-ET, the elastic force resulting from core compression and the radial surface tension counterbalancing it.

### Mechanical properties of the podosome

Knowing the elastic force, and thus the pressure *P*, allowed us to estimate the Young’s modulus of the core: *Y = P/ϵ_core_*= 1.7 MPa. While this value is several orders of magnitude larger than reported values for reconstituted and cellular actin networks ^29,30^, it is compatible with the very high actin density in the core. Actin itself has a Young’s modulus *Y_a_*= 2.3 GPa ^31^. The elastic modulus that can be reached by an actin network can be estimated as *Y = Y_a_ϕ^2^*, with *ϕ* the volume fraction occupied by actin filaments ^32^, yielding an upper limit of 12 MPa in podosomes. Thus, the Young’s modulus we find for the podosome core is well within expected values and helps understand how a compressive strain of less than 5% translates into forces in the nanonewton range.

The protrusive force exerted by the core on the substrate is balanced by an opposing force transmitted by the radial filaments to the adhesion ring, which prevents the core from relaxing towards the cell interior. We can estimate the surface tension, *σ*, of the 2D meshwork of radial filaments that is required to balance the force 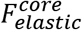, knowing the mean radius of the core, *r_core_*, and the mean angle of the radial filaments, *θ_radial_*, with respect to the membrane plane (Figure 4C and Figure S4B):

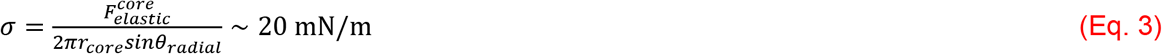

While this tension is one order of magnitude larger than the cortical tension of rounding mitotic cells ^33^, it is the same order of magnitude as the tension inferred for some adhering cells, from 6 mN/m ^34^ to 100 mN/m ^35^.

## Discussion

In summary, our results showed that mechanical energy accumulates inside the podosome core through bending of the dense network of actin filaments (Figure 1 and Figure 3). To relax, the elastic energy produces an elastic force, which is balanced by radial tension between the actin core and the adhesion ring. Indeed, we found that the elastic force computed from the bending energy matches the protrusive force measured experimentally (Figure 4). We revealed that the load borne by actin filaments in contact with the membrane is greater than the stall force, making their growth thermodynamically unfavorable (Figure 2). In addition, the need to synchronize their growth against the membrane would drastically slow down the polymerization process ^36^. Thus, how the network assembles under force remains to be understood.

One possibility is that actin filaments do not grow directly against the load at the membrane, which would instead be borne by the existing dense and compressed network. They would rather grow bent in the bulk of the podosome core in a dense environment, thus increasing the elastic energy stored by the network, which can then relax by pushing against the membrane. An estimation of the energy released by actin polymerization in the core, by summing over all actin monomers ^17^, yields 1-3.10^5^*k_B_T*, which is one order of magnitude larger than the elastic energy 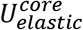. While some of the polymerization energy will be dissipated as heat, the polymerization reaction can still significantly contribute to the loading of the network.

Another possibility is that the filaments grow under a much smaller force, and that the network is then loaded by the active tension of the actomyosin cables driven by myosin II. This would be consistent with the decrease in pushing force when myosin II is inhibited ^24^. In this scenario, the energy source would be mostly ATP consumed by myosin II rather than the polymerization energy. We expect that a combination of experiments and simulation will be able to clarify the assembly of such a spring-loaded network.

In conclusion, to understand the forces applied by actin assemblies, it is not sufficient to consider only the local actin-membrane interactions; rather, the whole system must be taken into account. Through a combination of quantitative analysis and modelling of the native 3D architecture of macrophage podosomes, we showed that elastic energy is stored in actin networks *in vivo*, allowing forces of the order of ten nN to be produced. These results highlight the possibilities opened by exploring the architecture of actin networks *in situ* at the molecular scale.

Given the rapid progress of cryo-ET, we expect that, building on our biophysical approach, it will soon be possible to address other degrees of freedom for elastic energy storage, such as crosslinker deformation and filament twisting, and to shed light on other modes of force generation by cellular machineries.

## Methods

### Differentiation and culture of primary monocyte-derived macrophages

Human monocytes were isolated from the blood of healthy donors as described previously ^37^. Cells were resuspended in cold phosphate buffered saline (PBS) supplemented with 2 mM EDTA, 0.5% heat-inactivated Fetal Calf Serum (FCS) at pH 7.4 and magnetically sorted with magnetic microbeads coated with antibodies directed against CD14 (Miltenyi Biotec). Monocytes were then seeded on glass coverslips at 1.5 × 10^6^ cells/well in six-well plates in RPMI 1640 (Invitrogen) without FCS. After 2h at 37°C in a humidified 5% CO2 atmosphere, the medium was replaced by RPMI containing 10% FCS and 20 ng/mL of Macrophage Colony-Stimulating Factor (M-CSF) (Peprotech). For experiments, cells were harvested at day 7 using trypsin-EDTA (Fisher Scientific) and centrifugation (320g, 10 min).

### Cell vitrification

Gold EM grids with R1/4 holey SiO_2_ film (Quantifoil) were glow-discharged in a EasiGlow (Pelco) glow discharge system. After grid sterilization under UV light, the cell suspension containing fiducials was seeded onto the grids and incubated for 2 h at 37°C to let the cells adhere to the grids, resulting in 3 to 4 cells per grid square. For cell vitrification, grids were loaded into the thermostatic chamber of a Leica EM-GP automatic plunge freezer, set at 20°C and 95% humidity. Excess solution was blotted away for 10 s with a Whatman filter paper n°1, and the grids were immediately flash frozen in liquid ethane cooled at −185°C.

### Unroofing

When indicated, macrophages plated on grids were unroofed prior to vitrification. Cells were unroofed using distilled water containing complete™ protease inhibitors (Roche) and 10 μg/mL phalloidin (Sigma-Aldrich P2141) for 30 s.

### Cryo-FIB milling

Plunge-frozen EM grids were clipped into Autogrid frames modified for wedge milling under shallow angles ^38^. Autogrids were mounted into a custom-built FIB-shuttle and transferred using a cryo-transfer system (PP3000T, Quorum) to the cryo-stage of a dual-beam Quanta 3D FIB/SEM (Thermo Fisher Scientific) operated at liquid nitrogen temperature ^39^. The support film close to the cells of interest was sputtered away with high beam currents of 0.5-1.0 nA to provide a reference in Z direction for wedge milling. Cells were first milled roughly at very shallow angles (typically 2-5° of the incident ion beam) with beam currents of 300-500 pA. Advancement of the milling was monitored by SEM at 5 kV and 5.92 pA. Closer to the cell surface, beam currents of 50-100 pA were used for fine milling. Once all the wedges were prepared on the grid, a final polishing step at 30-50 pA was performed to limit surface contamination.

### Cryo-ET and tomogram reconstruction

Wedges were loaded vertically to the tilt axis in a Titan Krios transmission electron microscope (Thermo Fisher Scientific) equipped with a 300 kV field-emission gun, Volta phase plates (VPPs) (Danev et al., 2014), a post-column energy filter (Gatan, Pleasanton, CA, USA) and a 4k x 4k K2 Summit direct electron detector (Gatan) operated with SerialEM. The VPPs were aligned and used as described previously ^40^. Low-magnification images were recorded at 2250x. High-magnification tilt-series were recorded at 33,000x (calibrated pixel size 0.421 nm) with a target defocus for phase-plate imaging of 0 μm. Bi-directional tilt series were acquired typically from - 30° to +60° and −32° to −60° with a tilt increment of 2° and a total dose between 150 and 200 e-/Å^2^. Frames were aligned with in-house software (K2Align) based on procedures developed by Li et al. ^41^. Tilt series were aligned using the gold beads deposited on the surface of the support film as fiducial markers. 3D reconstructions with final pixel sizes of 1.684 nm were obtained by weighted-back projection using the IMOD software ^42^.

### Automated filament segmentation

The 4 x binned tomograms (pixel size of 1.684 nm) were subjected to nonlocal-means filtering using the Amira software ^43^ provided by Thermo Fisher Scientific. Actin filaments were traced using an automated segmentation algorithm based on a generic filament as a template ^44^, with a diameter of 8 nm and a length of 42 nm. To reduce background noise, short filamentous structures with lengths below 60 nm (or 50 nm for the unroofed podosomes) were filtered out.

The coordinates of the segmented filaments were exported from Amira and used as input for data analysis in MATLAB (The MathWorks) and in python (https://gitlab.com/jhervy/podosome-demo). The coordinates of the filaments were resampled every 3 nm to give the same weight to every point along a filament ^21^.

### Radial distance analysis

All the parameters (filament length, density and orientation, compressive strain and density of elastic energy) were evaluated as detailed in Figures S1–S3, S7 and S8, respectively. The plots as a function of the radial distance were computed by considering all the values in a given interval [*r,r*+δ] at a radial distance *r* from the core center. A list of 21 linearly-spaced values starting from 0 to 500 nm was used for the binning of the radial distance, which corresponds to an interval value of δ=25 nm (Figure 1D-F, Figure 3D-E, Figure S5A).

### Estimation of the core radius

All the parameters except the compressive strain were fitted as a function of the radial distance *r* from the core center using the following equation:

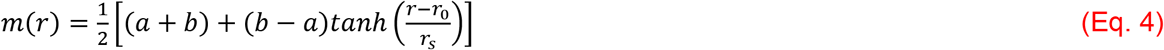

*m(r)* is either a decreasing function (for the filament density, filament orientation and the density of elastic energy) or an increasing function (for the filament length) with two saturation values set by the parameters *a* and *b* (Figure 1D-F and Figure 3E). The parameter *r_0_* corresponds to the radial distance for which the slope of the tangent line is maximum; the parameter *r_s_* defines the transition range between the two saturating values (Figure S5A). We used the parameter *r_0_* from the fit of the orientation as a measure of the core radius (Figure S5B).

### Polymerization force

To estimate the polymerization force generated at the core membrane, the protrusive filaments, *i.e*. those in the immediate vicinity of the plasma membrane, were considered. Specifically, all the filament portions up to 10 nm away from the membrane were taken into account (Figure 2A). A single filament growing perpendicularly to the membrane generates a force given by ^4^:

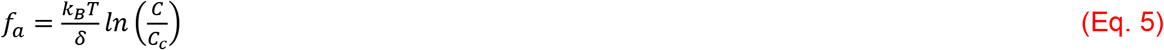

where *k_B_T*= 4.11 10^-21^ J, *δ* = 2.75 nm is the displacement induced by the addition of one actin monomer, *C* is the concentration of actin monomers in solution and *C_c_* is the critical concentration. Using the values *C* = 150 μM as estimated in ^45^ and *C_c_*= 0.06 μM for the critical concentration at the plus end as measured *in vitro*^46,47^, we found an upper limit of 11 pN for the stall force. Thus, the factor *ln(C/C_c_)* was set to 7 leading to a stall force of 10 pN for a single filament growing perpendicularly to the membrane. This value was increased by a factor *1/sin(θ)* for a filament having an orientation *θ* relative to the membrane ^17^ (Figure 2C). Therefore, the total polymerization force was computed as follows:

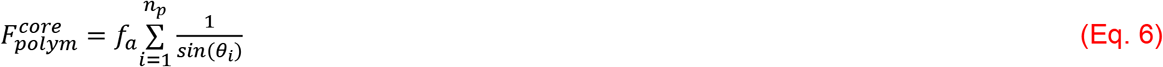

where *n_p_* is the number of protrusive filaments and *θ_i_* is the average orientation for the protrusive filament *i* (Figure 2D).

### Local curvature

The local curvature was evaluated between two consecutive points using their respective tangent vector *t* as follows:

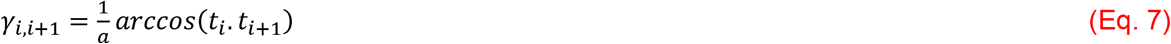

where *a* is their relative distance set to 3 nm in our segmentation procedure.

### Local elastic energy

The local energy is proportional to the square of the local curvature and can be discretized as follows:

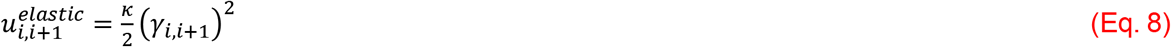

where *κ* is the bending modulus ^28^.

### Total elastic energy

The total elastic energy inside the core was computed by summing the local energy over all the filament points that are within the radial distance domain *r*∈[0,*r_core_*], where *r_core_* is the core radius (Figure S8).

### Elastic force

The elastic force pushing perpendicularly to the membrane is the derivative of the core elastic energy with respect to the height *h_core_* of the core:

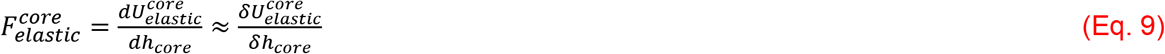

with *δ* indicating small changes. Indeed, in the core, the average compressive strain of the filaments is *ϵ_core_* ~ 0.04 (Figure 3D) and therefore considered to be small. Assuming all elastic energy to be released in the resting state, we have 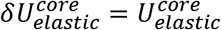 and *δh_core_ = ϵ_core_ × h_core_*.

Therefore, we find:

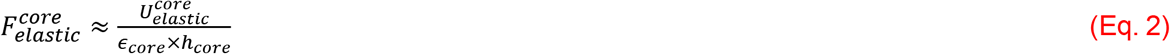

Note that this is an order of magnitude estimate: since the actin network is assembled under pressure, it is likely that its resting state has a non-zero elastic energy, in which case the force is overestimated. The precise architecture of the network could also yield a prefactor in the relation between *δh_core_* and *ϵ_core_*. Lastly, the elastic energy could be underestimated: part of it might be stored in crosslinker elasticity and in tension of the non-bent segments of actin filaments.

### Pressure induced in the podosome core

Assuming a circular shape of radius *r_core_* for the core, the pressure was computed as:

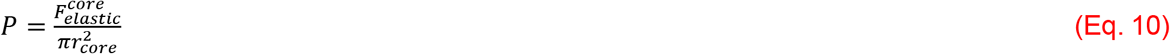

In the rest of this section, the values used to compute the pressure from the cryo-ET data and the PFM force measurements reported in ^16^ are detailed.

a. ***From the tomograms*** The pressure induced by the compression of the actin filaments (labelled as “Elastic”, Figure 4A) and by the polymerization at the membrane (labelled as “Polymerization”) were evaluated using the values for *r_core_* estimated from the fits of the orientation data (Figure 1E, inset).
b. ***From PFM force measurements*** The pressure *P_PFM_* was computed using the force value *F* = 10.4 ± 3.8 nN reported in ^16^ and *r_core_* = 203.1 ±37.8 nm estimated from the average of 10 podosomes (black dashed line in Figure 1E, inset). The error bar for this value was computed using the propagation of uncertainty method as follows:

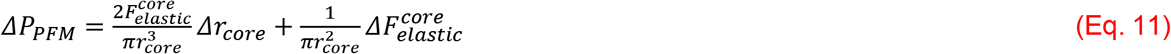 We found *P_PFM_*= 80.0 ± 59.0 kPa, with the margin of error *ΔP_PFM_* represented as a rectangular blue area filled in Figure 4B.

### Resource availability

All data and code used in the analysis are available upon request to the corresponding authors.

## Acknowledgements

This work benefited from the assistance of Vanessa Soldan from the Multiscale Electron Imaging platform (METi) of the Centre de Biologie Intégrative (CBI). The authors thank Martin Lenz for helpful discussions. This work was supported by the Human Frontier Science Program (RGP0035/2016), la Fondation pour la Recherche Médicale (FRM DEQ2016 0334894), a CNRS Momentum fellowship, l’Agence Nationale de la Recherche and Deutsche Forschungsgemeinschaft (ANR-DFG JA-3038/2-1) and with financial support from ITMO Cancer of Aviesan on funds managed by Inserm.

## Contributions

MJ, RP designed the project. AB, RP prepared the cells. SB vitrified the cells. MJ performed cryoFIB milling, cryo-ET, tomogram reconstruction and segmentation. MJ, JH, AP, RP designed and performed the experimental analysis. RV, SD designed the theoretical analysis. JH, SD performed the theoretical analysis. MJ, SD, RP supervised the project. MJ, IMP, WB, SD, RP obtained funding. MJ, JH, SD, RP wrote the manuscript with input from the others.

## Ethics declaration

The authors declare no competing interests.

## Supplemental information

**Movie S1. Native podosome architecture revealed by *in situ* cryo-ET.**

This movie shows the tomogram of the podosome presented in Figure 1A-C followed by its segmentation. Actin filaments are colored as a function of their orientation with respect to the basal membrane. Intermediate filaments are in brown and membranes in grey. Successive rotation, sectioning and zoom in the segmented volume allow the visualization of the dense organization of the actin filaments of the podosome core.

**Movie S2. Architecture of unroofed podosomes by cryo-ET.**

This movie shows the tomogram of unroofed podosomes presented in Figure S6 followed by its segmentation. Actin filaments are colored as a function of their orientation with respect to the basal membrane. Successive rotations, sectioning and zoom in the segmented volume reveal the nanoscale actin organization of neighboring podosomes.

**Movie S3. Visualization of the actin filaments in the vicinity of the plasma membrane beneath a podosome core.**

This movie shows the actin segmentation of the podosome presented in Figure 2A. Core filament segments in the vicinity of the basal membrane are colored as a function of their orientation. A series of rotations, zoomed and unzoomed sections through the segmented volume allow to visualize the number and orientation of the protrusive filaments at the core.

**Movie S4. Podosome actin filaments are bent.**

This movie shows the actin segmentation of the podosome presented in Figure 3A. Actin filaments are colored as a function of their mean curvature. Successive zoomed sections through the segmented volume allow to visualize the curvature of the filaments in a native podosome.

**Figure S1.**
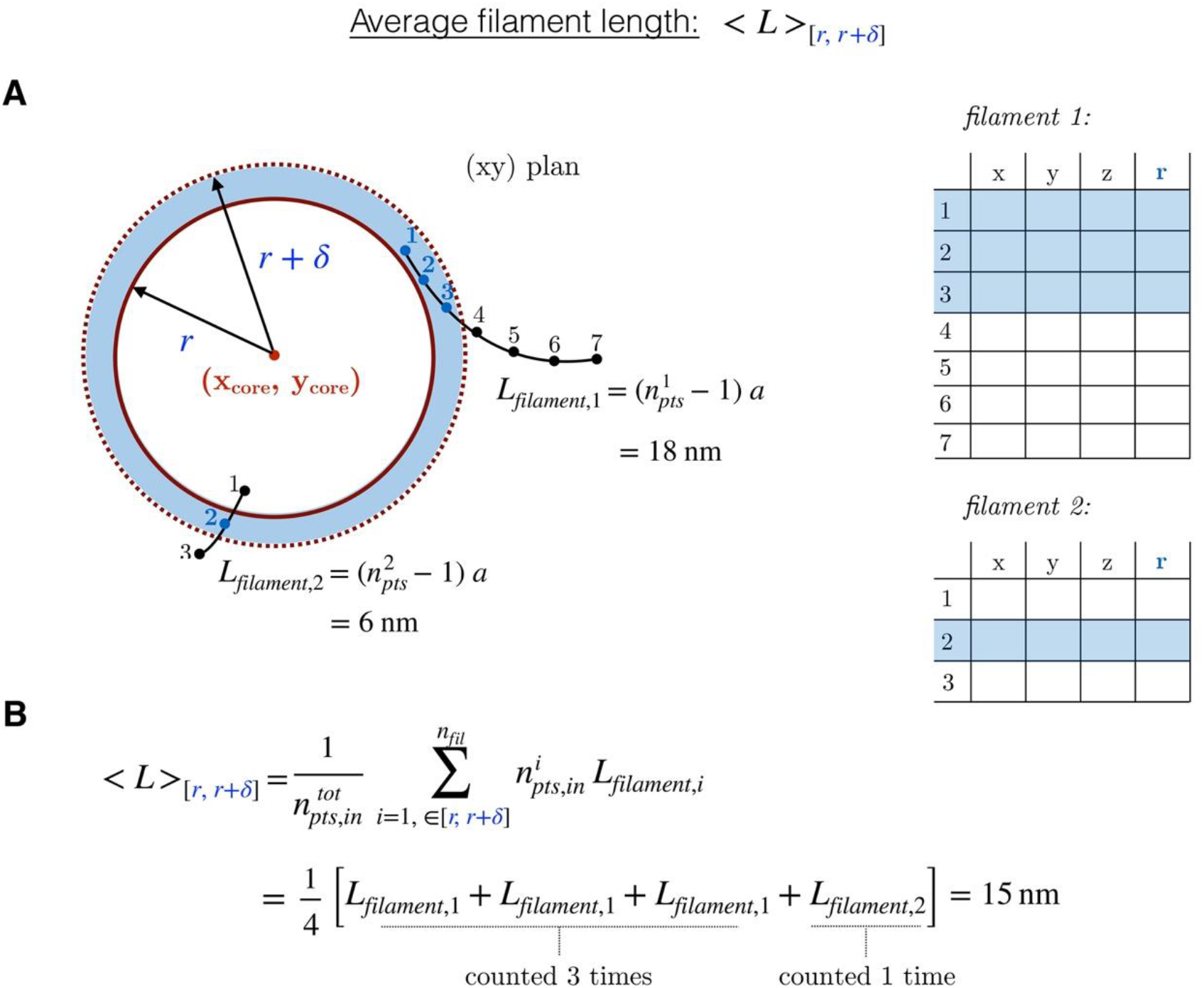
Illustration of the computation of the average filament length as a function of the radial distance. **A:** For each distance bin [r,r+δ], the points inside the bins (shown in blue) are identified for each actin filament (left). Each point is represented both in cartesian and radial coordinates (right). **B:** The average length in the bin is the mean of the filament lengths weighted by their number of points in the bin.

**Figure S2.**
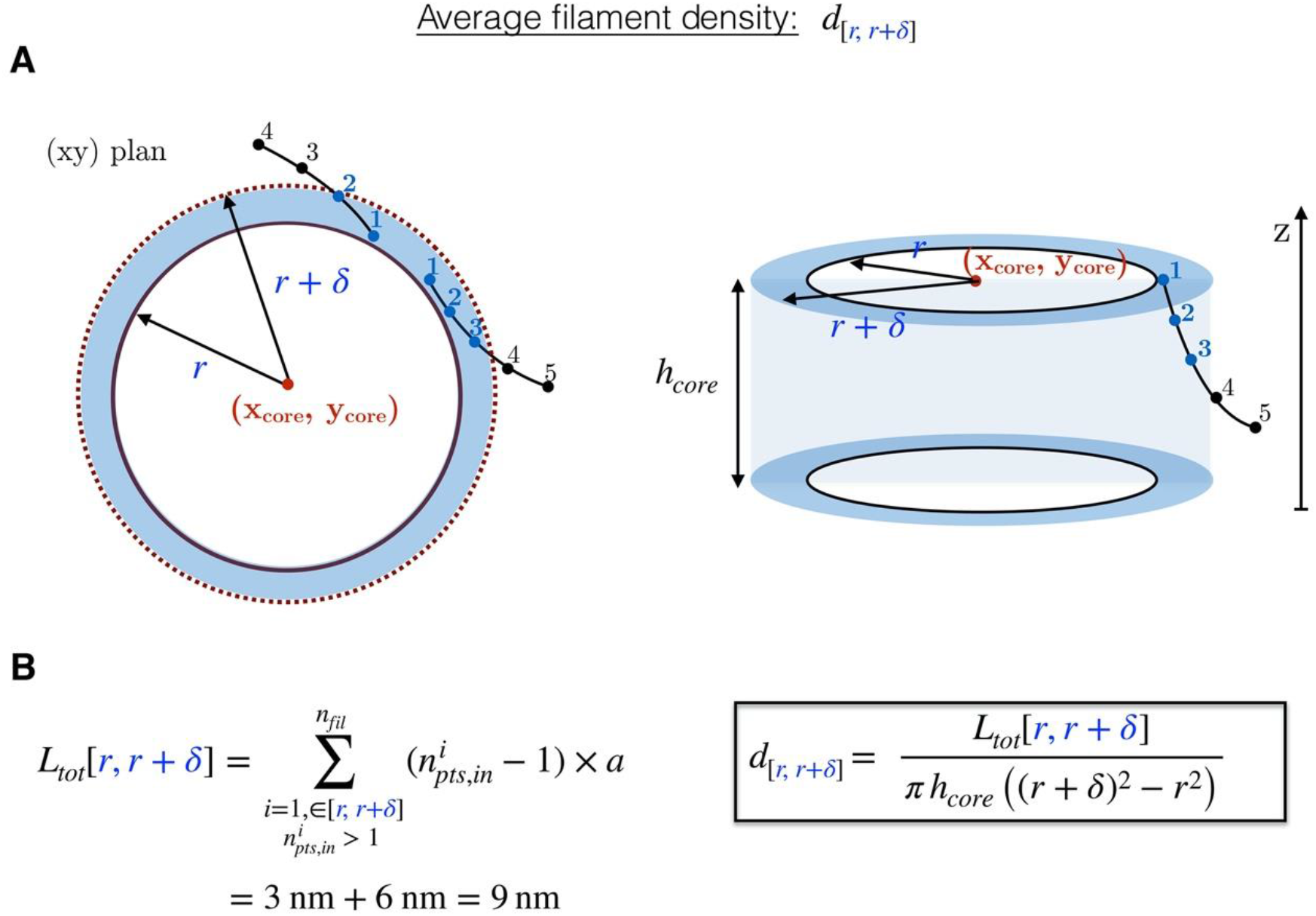
Illustration of the computation of the average filament density as a function of the radial distance. **A:** For each bin [r,r+δ], the points inside the bins (shown in blue) are identified for each actin filament. **B:** The total actin length is calculated in each bin and divided by the bin volume to compute the density.

**Figure S3.**
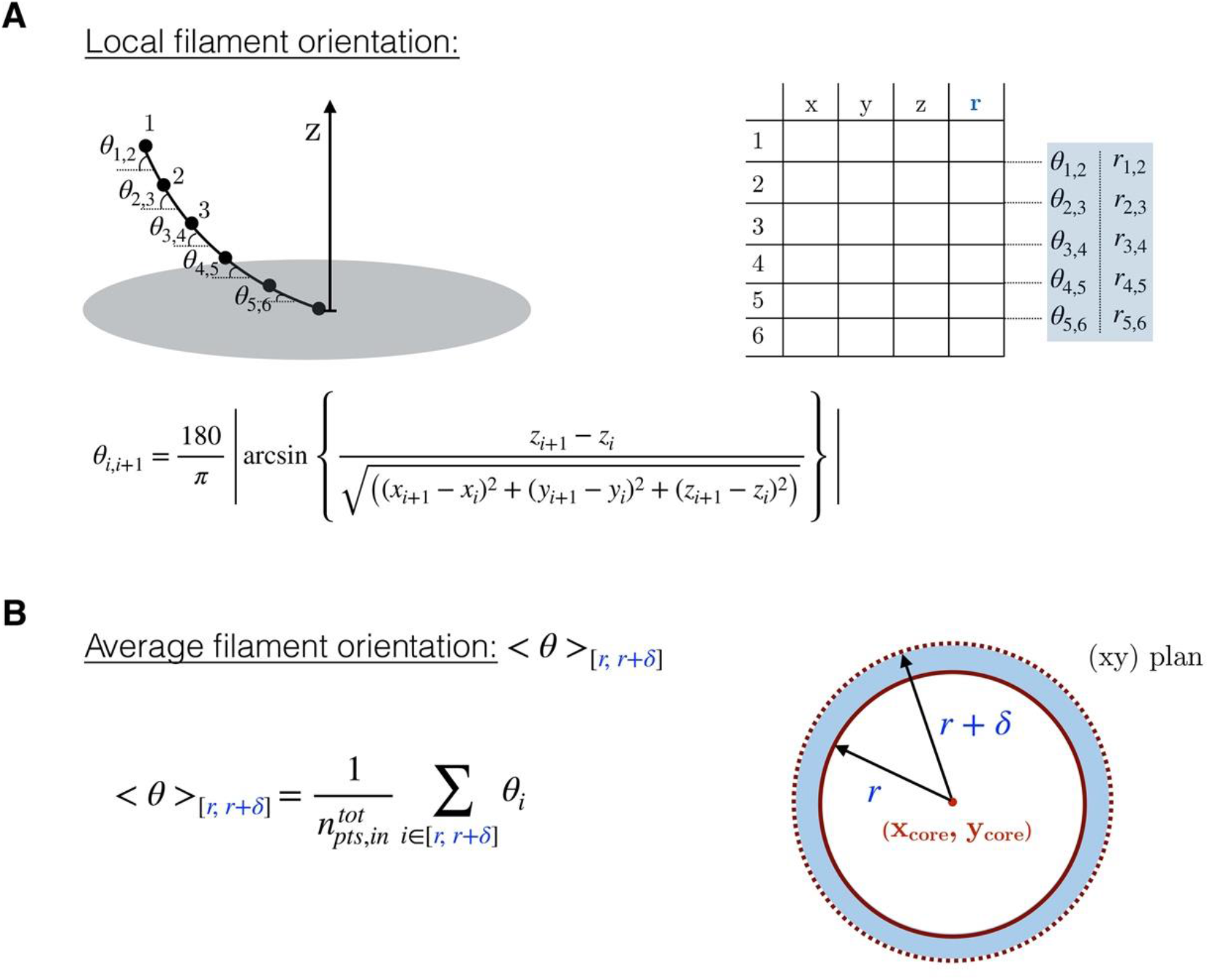
Illustration of the computation of the average filament orientation as a function of the radial distance. **A:** To each segment between two filament points is associated an angle with respect to the membrane plane. **B:** The average filament orientation in a bin is the mean of the orientations of all segments inside the bin.

**Figure S4.**
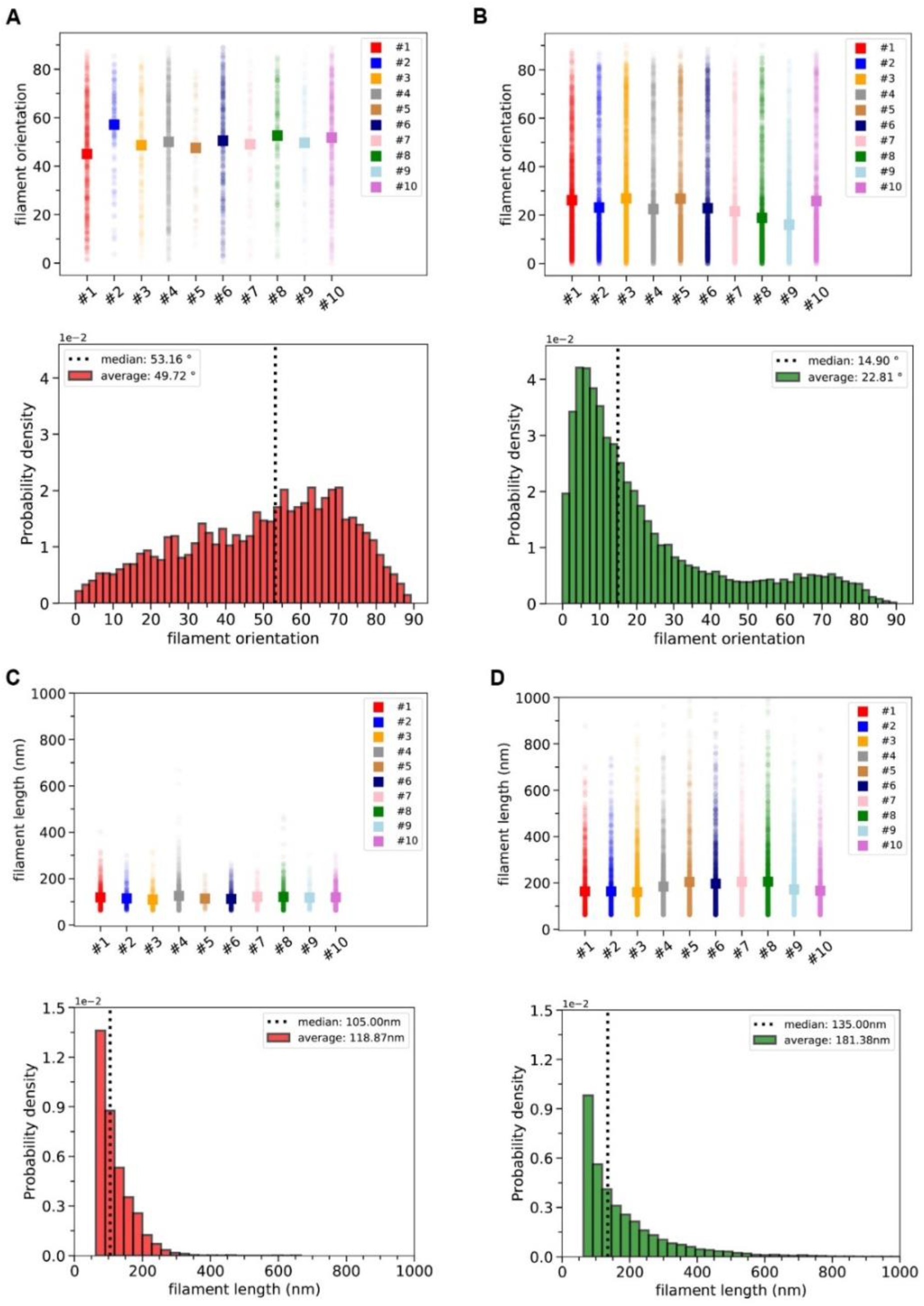
Orientation and length distributions of the innermost and outermost filaments, respectively, in ten podosomes. **A:** (Top) Mean orientation of the innermost filaments within the radial distance range rϵ [r_0_-r_s_,r_0_]. r_0_ and r_s_ are the inflection point and transition range parameters, respectively, and were obtained from the fit of the orientation as a function of r for each tomogram. (Bottom) Corresponding distribution obtained by merging all the data from the top panel into a single dataset. The vertical dashed line indicates the median value of the dataset. **B:** Same distributions for the outermost filaments, *i.e.* within the radial distance range r ϵ[r_0_+r_s_,r_max_] where r_max_ is the highest radial distance between a point filament and the core center. **C-D:** Same plots as in **A-B** for the filament length.

**Figure S5.**
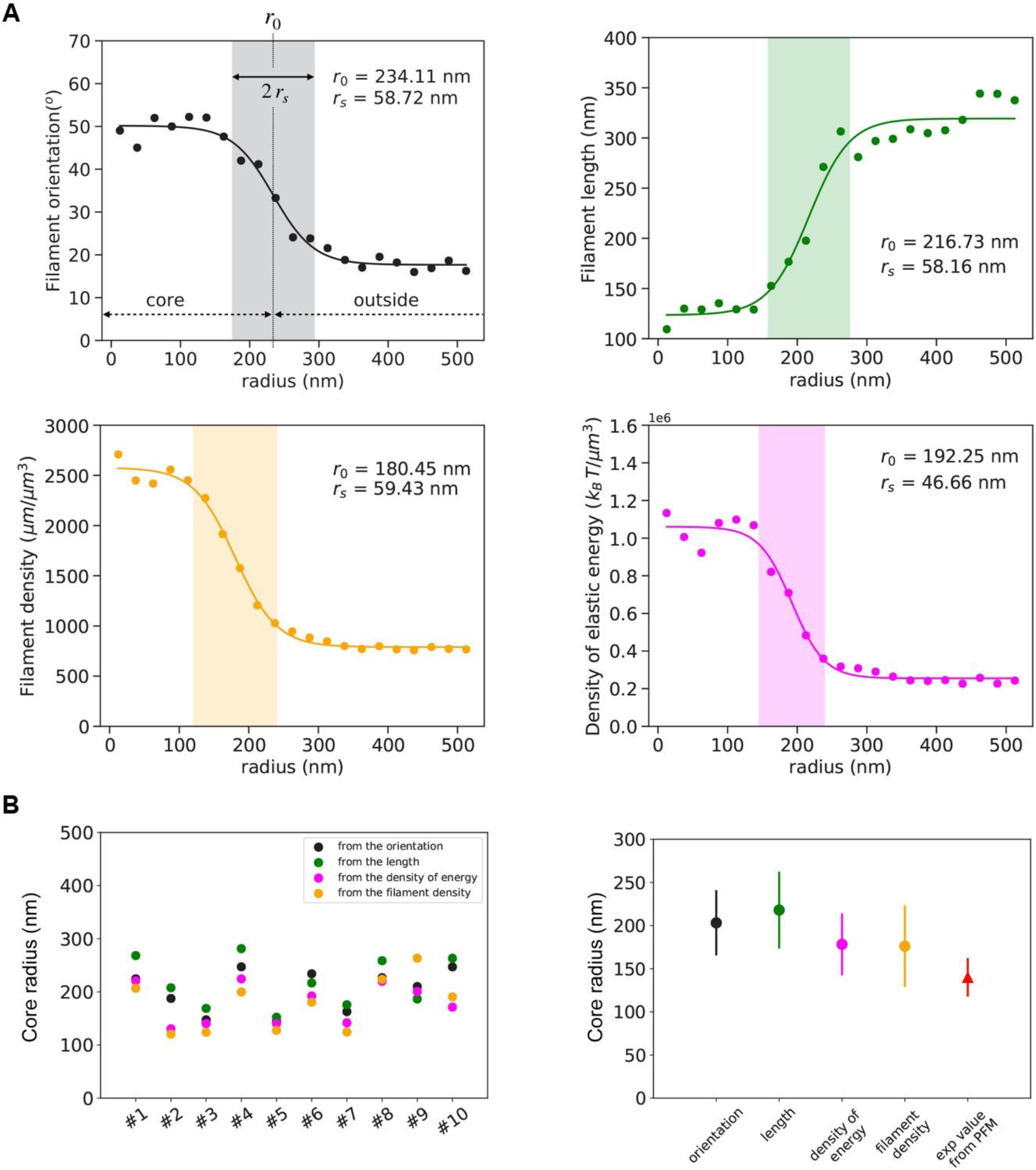
Estimation of the core radius from the distribution of each parameter as a function of the distance to the core center. **A:** Filament orientation, length, actin density and density of energy for podosome #6 with their respective fits are shown. The parameters r_0_ and r_s_ and the transition (colored) zone are indicated. **B:** (Left) Comparison of the core radius values obtained from the fits of the different parameters as a function of the radial distance for the ten tomograms. (Right) Average radius of the podosome core estimated from the quantitative analysis of the cryo-ET data and compared to the reported value from PFM (mean±s.d.).

**Figure S6.**
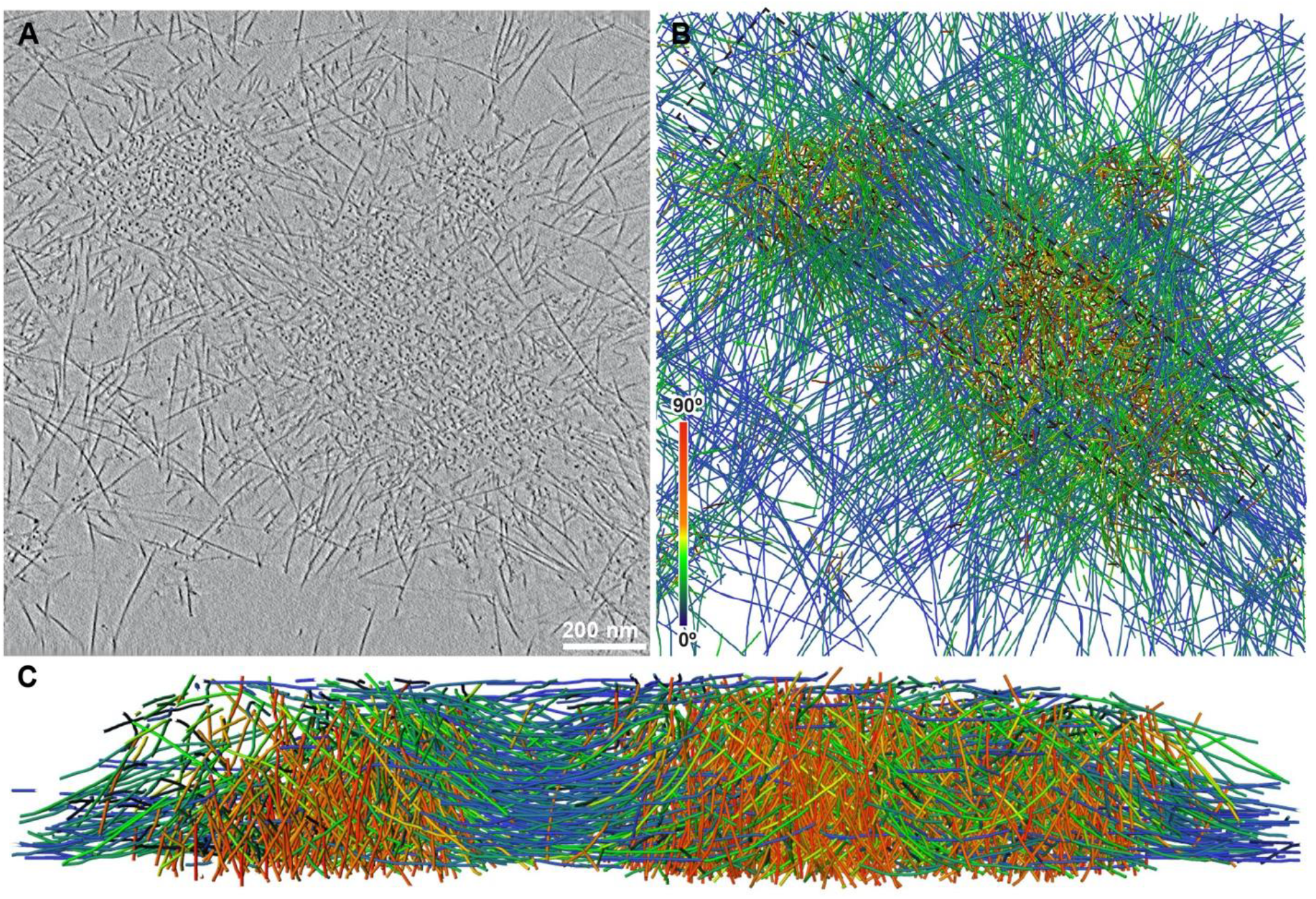
Unroofed podosome architecture revealed by cryo-ET. **A:** Slice from a tomographic volume acquired in a plunge-frozen, unroofed human macrophage revealing neighboring podosomes. **B:** Orthographic view of the corresponding 3D segmentation of the actin filaments showing their relative orientation with respect to the basal membrane. **C:** Perspective view of the volume indicated by a dotted rectangle in (D) and shown from the left side. See also Movie S2.

**Figure S7.**
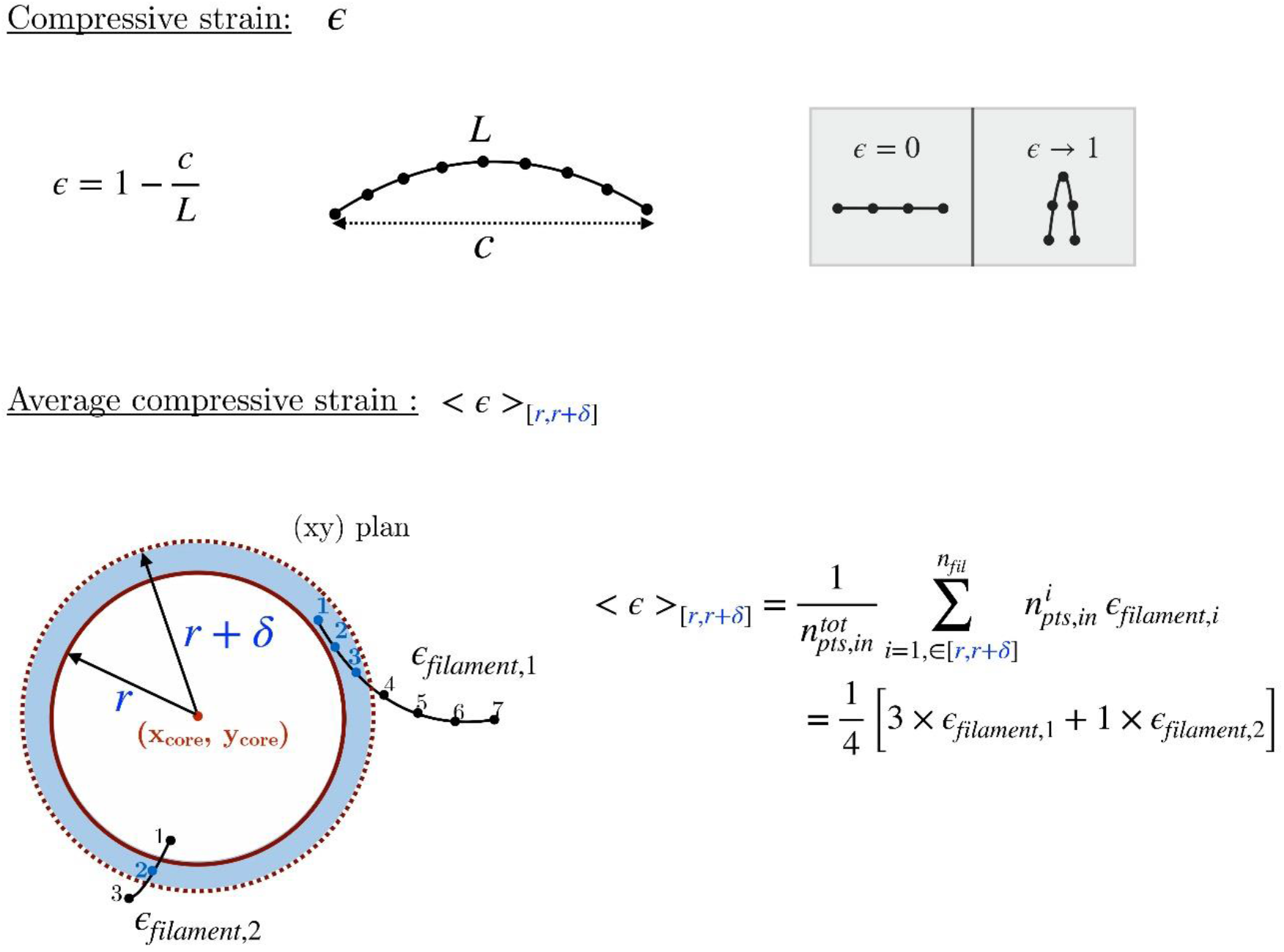
Illustration of the computation of the average compressive strain as a function of the radial distance. **A:** The compressive strain is computed as 1 minus the ratio between the end-to-end distance, *c*,and the filament length, *L*. Thus, a straight filament has a zero strain. **B:** The average compressive strain in a bin is computed as the mean of the compressive strains weighted by the number of filament points in the bin.

**Figure S8.**
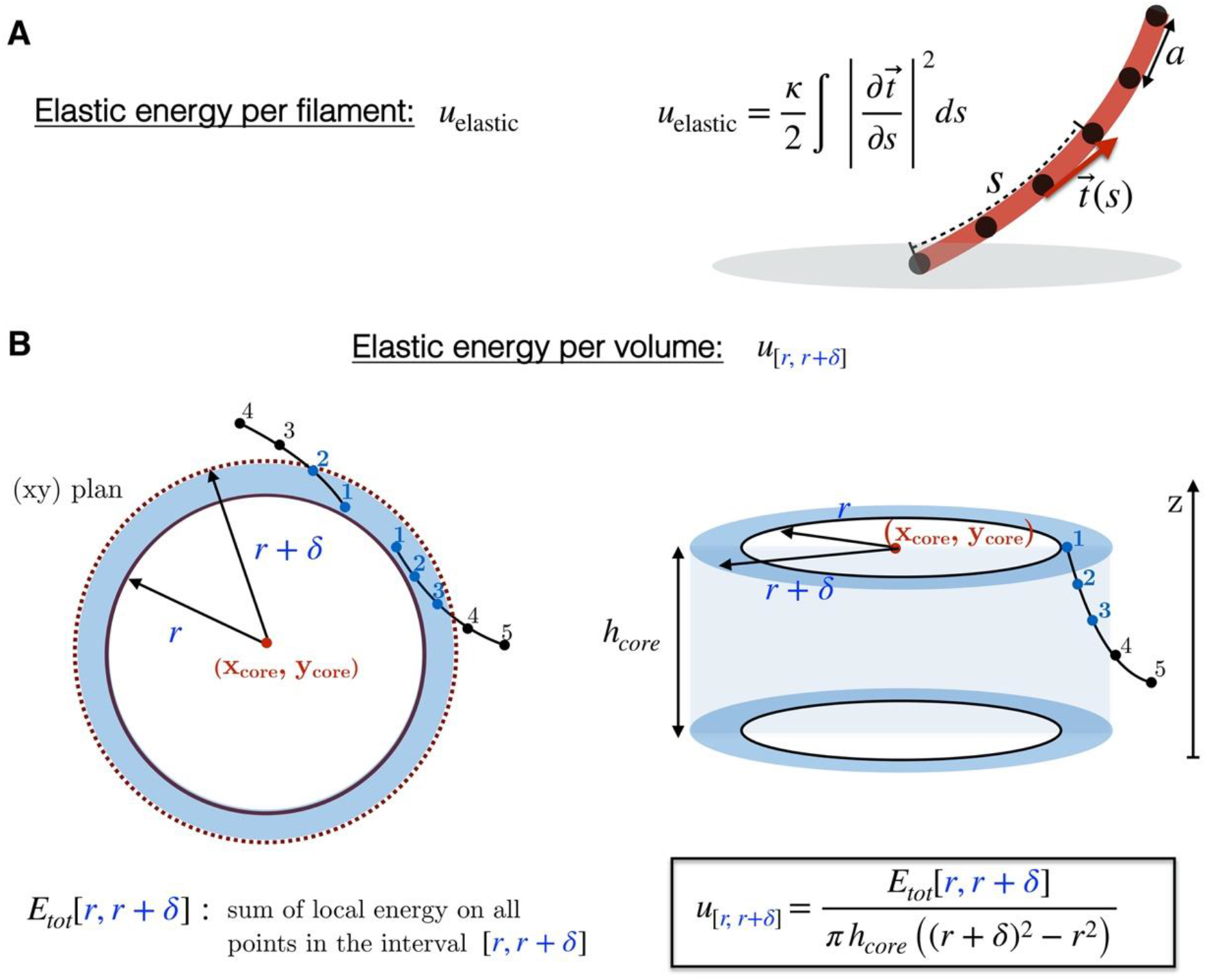
Illustration of the computation of the elastic energy per volume as a function of the radial distance. **A:** Definition of the elastic energy of a filament as the integral of the squared curvature over the filament length. **B:** The density of elastic energy as a function of the radial distance is computed as the sum of the elastic energy of all the filament points in the bin divided by the volume.

## Notes

### Competing Interest Statement

The authors have declared no competing interest.

